# Multi-modal brain fingerprinting: a manifold approximation based framework

**DOI:** 10.1101/209726

**Authors:** Kuldeep Kumar, Laurent Chauvin, Matthew Toews, Olivier Colliot, Christian Desrosiers

## Abstract

This work presents an efficient framework, based on manifold approximation, for generating brain fingerprints from multi-modal data. The proposed framework represents images as bags of local features, which are used to build a subject proximity graph. Compact fingerprints are obtained by projecting this graph in a low-dimensional manifold, using spectral embedding. Experiments using the T1/T2-weighted MRI, diffusion MRI, and resting state fMRI data of 945 Human Connectome Project subjects demonstrate the benefit of combining multiple modalities, with multi-modal fingerprints more discriminative than those generated from individual modalities. Results also highlight the link between fingerprint similarity and genetic proximity, monozygotic twins having more similar fingerprints than dizygotic or non-twin siblings. This link is also reflected in the differences of feature correspondences between twin/sibling pairs, occurring in major brain structures and across hemispheres. The robustness of the proposed framework to factors like image alignment and scan resolution, as well as the reproducibility of results on retest scans, suggest the potential of multi-modal brain fingerprinting for characterizing individuals in a large cohort analysis. In addition, taking inspiration from the computer vision community, the proposed rank retrieval evaluation based on the task of twin/sibling identification and using Mean Average Precision (MAP) can be used for a standardized comparison of future brain fingerprints.

## 1. Introduction

Despite sharing gross similarities, individual brains show a significant amount of variability [1] in terms of structure [2], function [3, 4, 5], and white matter architecture [6, 7]. Recently, various studies have focused on characterizing this
variability using brain *fingerprints*, for instance, based on shape [8], functional connectivity [9, 10], white matter fiber geometry [11] or voxel-wise diffusion density [12]. These studies are motivated by the fact that brain characteristics are largely determined by genetic factors that are often unique to individuals [13]. Moreover, various neurological disorders like Parkinson [14] and autism [15] have been linked to specific brain abnormalities that are difficult to describe at the population level. With the rapid improvements in MRI acquisition hardware and analysis tools, and thanks to large brain-related initiatives like the Human Connectome Project (HCP) [16] and UK Biobank [17], researchers are better poised to study individual subjects in addition to groups [18, 19], thus taking a step towards precision medicine [20] and precision psychiatry [21].

The importance of brain fingerprinting is evident from the recent surge in studies on this topic. For example, Yeh et al. [12] built a local connectome fingerprint using dMRI data, and applied this fingerprint to the analysis of genetically-related subjects. Kumar et al. [11] proposed another dMRI-based fingerprint called *Fiberprint*, which characterizes white matter fiber geometry. Finn et al. [9] considered the correlation between time courses of atlas-defined nodes to generate a functional connectivity profile, and used this profile to identify individuals across scan sessions, both for task and rest conditions. Liu et al. [10] use dynamic brain connectivity patterns for Identifying individuals and predicting higher cognitive functions. Moreover, Miranda et al. [22] proposed a linear model to describe the activity of brain regions in resting-state fMRI as a weighted sum of its functional neighboring regions. Their functional fingerprint, derived from the model's coefficients, has the ability to predict individuals using a limited number of non-sequential frames.

Various morphometry-based fingerprints have also been proposed for structural MRI modalities like T1- or T2-weighted images. For example, Wachinger et al. [8] quantify the shape of cortical and subcortical structures via the spectrum of the Laplace-Beltrami operator. The resulting representation, called *Brainprint*, is used for subject identification and analyzing potential genetic influences on brain morphology. Toews et al. [23] represent images as a collection of localized image descriptors, and apply scale-space theory to analyze their distribution at the characteristic scale of underlying anatomical structures. This representation is employed to identify distinctive anatomical patterns of genetically-related individuals or subjects with a known brain disease.

So far, fingerprinting studies in the literature have focused on a single modality. However, each modality captures unique properties of the brain and combining multiple modalities can provide a richer, more discriminative information [24, 25]. Hence, the fusion of multiple modalities has been shown superior than single-modality data to identify diseases like schizophrenia, bipolar disorder, major depressive disorder and obsessive-compulsive disorder [24]. Multi-modal neuroimaging biomarkers have also been proposed to predict cognitive deficits in schizophrenia [26]. Similarly, the combination of multiple MRI modalities has led to the improved segmentation of isointense infant brain images [27]. Also, multimodal imaging data can be used to predict the brain-age of subjects and detect cognitive impairments [28]. Detailed reviews on multi-modal methods and investigations for psychopathology can be found in [24, 29, 30].

Due to the challenges of combining multiple modalities in a single framework [24, 30], defining a multi-modal brain fingerprinting remains to this day an elusive task. Morphometry-based approaches, such as Brainprint [8], could potentially be extended to other modalities like dMRI. However, this requires solving non-trivial problems such as the cross-modality alignment of images with different resolutions, the segmentation and correspondence of neuroanatomical structures, etc. Computational efficiency is another important issue when dealing with large-scale, multi-subject and multi-modal datasets like the Human Connectome Project (HCP) [16] and UK Biobank [17]. In this work, we propose a multi-modal brain fingerprinting that overcomes these challenges using manifold approximation. The key idea is to represent each image as a bag of local features, and derive a subject-level proximity graph using feature correspondences over the entire set of images [23]. This subject proximity graph provides an approximation of the image appearance subspace (i.e., the manifold), which can be used to obtain a compact fingerprint representation.

Manifold learning has been extensively studied in machine learning [31] with many approaches like Isomap [32], Locally Linear Embedding (LLE) [33], Spectral Embedding [34] and Multi-dimensional Scaling (MDS) [35] proposed over the years. As detailed in [36], such techniques have also been used for various problems of medical imaging like registration, segmentation and classification. For example, in [37], Gerber et al. use manifold learning to perform a population analysis of brain images. Similarly, a deep learning based approach is explored in [38] to learn the manifold of brain MRIs. A key factor in such methods is image representation. For instance, the manifold could be approximated using the Euclidean distance between image pairs, however this would not be robust to translation, rotation or scaling, and would suffer from high computational costs. Representations based on local features, often referred to as bag of features (BoF) representations, have been shown to automatically identify known structural differences between healthy controls and Alzheimer’s subjects in a fully data driven fashion [23]. The ability to identify anatomical patterns that may only be present in subsets of subjects and without the stringent requirement of one-to-one correspondence between subjects, the BoF approach is well suited to capture disease or anatomical variability and to carry out large scale analysis.

BoF representations play a key role in various problems of computer vision, for example, object recognition [39, 40] and image retrieval [41]. This technique can be seen as a way of compressing full images using a few discriminative local features, which can then be matched in sublinear time, for example, using randomized KD-search trees [42]. With respect to brain imaging, BoFs have been used for morphometry analysis [23], modeling the development of infant brains [43], and image alignment [44]. While they have shown great potential for computer vision and medical imaging, BoFs have, thus far, not been explored for brain fingerprinting. This is mainly due to the fact that BoF representations can have a large and variable size, which makes comparing two fingerprints non-trivial. In this work, this problem is circumvented by embedding the BoF representations in a low-dimensional manifold.

The key contributions of this work are:

1. **Novel framework:** A novel and data driven approach based on Bag of Features (BoFs) and manifold approximation that combines the information from multiple imaging modalities into a single compact fingerprint;
2. **Large scale analysis:** Comprehensive analysis using pre-processed data from Human Connectome Project for T1/T2-weighted MRI, diffusion MRI (DTI and GQI measures), and resting state fMRI (netmats).
3. **Modality comparisons:** Quantifying contribution of individual modalities for fingerprint and validation of hypothesis that each modality provides certain complimentary information, using a common task of twin/sibling identification.

In addition, the current study presents a comprehensive analysis of the proposed fingerprint using a large-scale dataset from the Human Connectome Project (HCP), where numerous properties/factors are investigated, including the impact of various fingerprint parameters (e.g., manifold dimensionality and proximity graph connectivity), the contribution of individual modalities and/or their combination to the fingerprint’s discriminativeness, the fingerprint’s robustness to image alignment and scan resolution, and the reproducibility of results using re-test or corrupted scans.

Using the genetically verified Zygosity labels from the HCP twin dataset, we analyze the proposed fingerprint’s ability to identify genetically-related subjects (i.e., monozygotic twins, dizygotic twins and non-twin siblings) from a large cohort, and show our multi-modal fingerprint to outperform single-modality approaches or fingerprints based on raw images. In an analysis of local feature correspondences, we identify for each modality the neuroanatomical regions having the most significant differences across groups of genetically-related subjects, as well as between males and females. Lateral asymmetry is also considered in this analysis by comparing the distribution of features correspondences across hemispheres. To our knowledge, this study constitutes the most in-depth investigation of a multi-modal brain fingerprint.

This work extends our preliminary work [45, 46] in terms of 1) set of multiple modalities and recent HCP data release 2) manifold approximation and compact fingerprint generation 3) the rank retrieval analysis using Mean Average Precision and impact of various factors. Taking inspiration from computer vision challenges, the proposed rank retrieval evaluation based on the task of twin/sibling identification and using Mean Average Precision (MAP) can be used for a standardized comparison of future brain fingerprints. Also, twin data from Human Connectome Project (Q3 release) has been analyzed in various studies including an extensive analysis of heritability of multi-modal functional connectivity in [47]. Our work, compliments these studies as well as previous fingerprint studies in terms of the number of modalities used in a single analysis: structural MRI, diffusion MRI, resting state functional connectivity profiles, and their combinations. The proposed framework is computationally efficient and validates (quantitatively) the hypothesis that individual modalities provide certain complimentary information. In addition, while the genetic basis of brain structure and function is largely unknown [48], the neuro-anatomical traits are largely heritable [49, 50, 13] and form the basis for the identification of twins. The scope of the present study is limited to multi-modal brain fingerprinting, and a heritability analysis on the lines of Ge et al. [49] will be assessed in next step.

The rest of this paper is organized as follows. We first present the proposed multi-modal brain fingerprinting framework, detailing the data pre-processing steps, the BoF representation and proximity graph computation, and the manifold embedding of this graph. In Section 3, we then conduct an extensive experimental validation using the T1-weighted, T2-weighted, diffusion-weighted MRI, and resting state fMRI data of 945 subjects from the HCP dataset. Finally, we conclude with a summary of our contributions and a discussion of possible extensions.

## 2. Materials and methods

Figure 1 summarizes the pipeline of the proposed multi-modal brain fingeprint framework, which is comprised of four steps. In the first step, we start with pre-processed structural MRI (sMRI) and diffusion MRI (dMRI) data of 945 subjects from the Human Connectome Project [51, 16]. Diffusion Tensor Imaging (DTI) and Generalized Q-Ball Imaging (GQI) based Diffusivity measures are obtained from dMRI scans, including: fractional anisotropy (FA), axial diffusivity (AD), mean diffusivity (MD), radial diffusivity (RD) and generalized fractional anisotropy (GFA). The second step then extracts local features from the images of each subject, and encodes subjects as a bag of features (BoF). In the third step, the multi-modal fingerprints of subjects are computed using manifold approximation. Towards this goal, a subject-level proximity graph is first constructed by matching the features of each modality across images, and identifying pairs of subjects with a high number of correspondences. Fingerprints are then obtained by embedding this graph in a low-dimensional subspace. In the last step, we perform various analyses on the subject fingerprints. The informativeness of individual modalities and their link to genetic proximity are first measured in a twin/sibling identification analysis. This analysis is then extended to multi-modal fingerprints, showing the combined effect and complementarity of multiple modalities. Resting state fMRI network matrices and FreeSurfer derived measures of volume, thickness, and area, provided by HCP, are also used for fingerprint analysis. Finally, the distribution of feature correspondences between pairs of subjects are used to identify regions showing significant differences across different sibling types. The following subsections describe each of these steps in greater detail.

**Figure 1:**
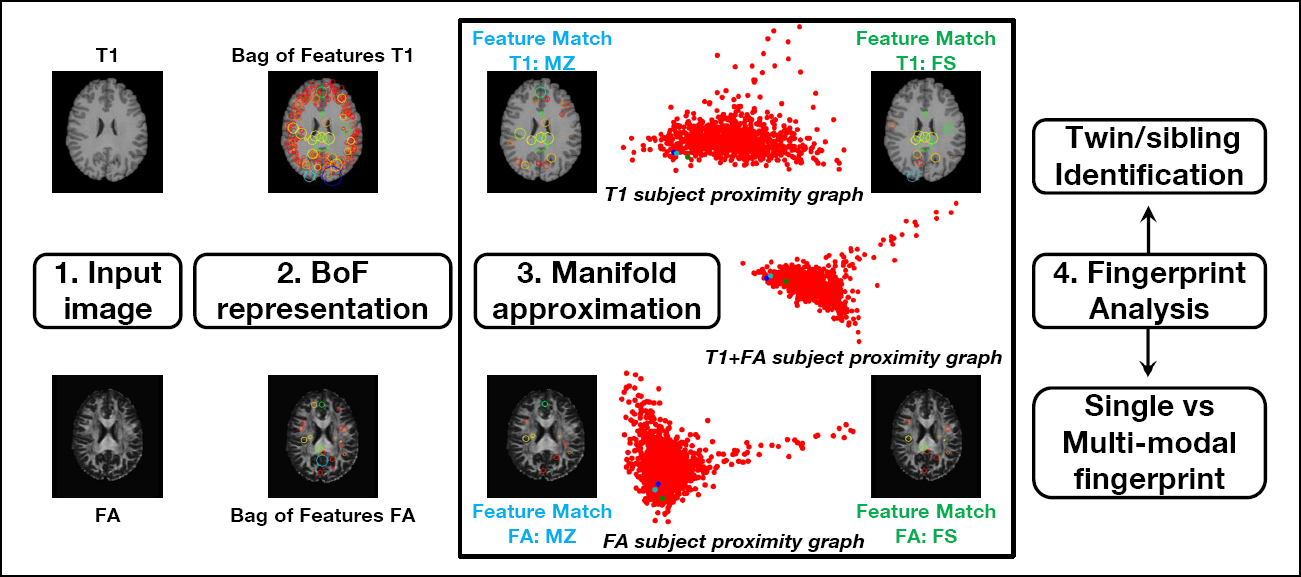
Pipeline of the proposed framework and analysis. For a given input image, a BoF representation is first obtained by extracting local features. This representation is then converted to a fingerprint by matching features across the entire set of images, and embedding the resulting proximity graph into the manifold. The manifold approximation block shows the 2D embedding coordinates (i.e., fingerprint) of HCP subjects (red dots) obtained with T1w (top), FA (bottom) and combined T1w+FA (middle) images. The fingerprints of a specific subject (blue dot), his/her monozygotic twin (MZ, cyan dot) and full sibling (FS, green dot) are highlighted in each plot. The pairwise feature matches of these two sibling pairs, for T1w and FA images, are shown in the corner images of the block.

### 2.1. Data and pre-processing

We used the pre-processed structural and diffusion MRI data, and resting state fMRI network matrices of 945 subjects, the retest data of 42 subjects, from the HCP1200 release of the Human Connectome Project [16]. The HCP1200 release provides genetically verified labels for twins and siblings, and is a rich resource to analyze the importance of environmental and genetic influences for traits, phenotypes, and disorders [52, 51]. Table 1 provides the demographic details of the subjects used in this study.

**Table 1:** Demographics. We considered the HCP1200 release subjects with structural MRI, diffusion MRI, and resting state fMRI netmats data, and for which the HasGT field is true (genetically verified data). The family structure and links are obtained from the output of hcp2blocks.m script listed in data release manual. The sibship sizes are between 1 and 6.

| Type | Total | Gender |  | Age | Handedness |  |
| --- | --- | --- | --- | --- | --- | --- |
|  |  | F | M | Range (median) | L | R |
| All | 945 | 501 | 444 | 22-36 (29) | 84 | 861 |
| MZ | 238 | 138 | 100 | 22-36 (30) | 19 | 219 |
| DZ | 126 | 70 | 56 | 22-35 (29) | 13 | 113 |
| NotTwin | 581 | 293 | 288 | 22-36 (28) | 52 | 529 |

Data were acquired on a Siemens Skyra 3T scanner [53] and pre-processed as described in [54]. The structural acquisitions include high resolution T1-weighted (T1w) and T2-weighted (T2w) images (0.7 mm isotropic, FOV = 224 mm, matrix = 320, 256 sagittal slices in a single slab), the diffusion acquisition used following parameters: sequence = Spin-echo EPI; repetition time (TR) = 5520 ms; echo time (TE) = 89.5 ms; resolution = 1.25 × 1.25 × 1.25 mm^3^ voxels, and the resting-state fMRI acquisition involved four 15min runs at 2 mm isotropic resolution and a repetition time of 0.72s (4800vol per subject). Further details can be obtained from the HCP1200 data release manual^2^. We used the `hcp2blocks.m` script (described in the HCP1200 release) to generate a FamilyID based matrix, only considering subjects having dMRI, sMRI, and rfMRI netmats data, and for which the `HasGT` field is true. Using this selection criterion, we obtained a total of 238 monozygotic (MZ) subjects, 126 dizygotic (DZ) subjects, and 581 non-twin subjects. The sibship size ranged between 1 and 6. In a next step, using the mother ID, father ID, family ID and family type information from the output of `hcp2blocks.m` script, we obtained 119 monozygotic twin pairs, 63 dizygotic twin pairs, 546 full-sibling (FS) pairs, 39 maternal half sibling (MHS) pairs, and 5 paternal half sibling (PHS) pairs. These pairs are used for twin/sibling identification task in the following sections.

For structural MRI we considered T1-weighted (0.7 mm) and T2-weighted (0.7 mm), with and without skull. The images are in native space and skull stripped, unless explicitly specified. In the case of dMRI data, signal reconstruction was performed with the freely available DSI Studio toolbox [55] using the Diffusion Tensor Imaging (DTI) and Generalized Q-Ball Imaging (GQI) reconstruction options. Four widely used DTI based measures were extracted to characterize white matter micro-structure: fractional anisotropy (FA), axial diffusivity (AD), mean diffusivity (MD) and radial diffusivity (RD). The interpretation of these measures are discussed in [56]. In addition, to utilize the multi-shell information and high angular resolution of the HCP data, Generalized Q-Ball Imaging (GQI) [55] based measures including generalized fractional anisotropy (GFA) and Quantitative Anisotropy (QA) were also obtained. For resting state fMRI, we used the connectivity matrices (netmats), provided by the HCP 1200 release, derived using the FSLNets toolbox, either via full correlation or the partial correlation [57], the latter being calculated by inverting the covariance matrix. For analyzing the impact of alignment, we also used the MNI space aligned data for T1-weighted (0.7 mm) and T2-weighted (0.7 mm) provided by the HCP 1200 release. In addition, to combine structural modalities with dMRI, and to analyze impact of scan resolution, we re-sampled T1- and T2-weighted images to a 1.25 mm resolution, using the linear registration (FLIRT) tool of FSL [58]. Our analysis also utilized FreeSurfer derived measures of sub-cortical volumes, cortical thickness and area, as well as T1w/T2w MRI ratio images (0.7 mm, myelin content information).

### 2.2. Multi-modal brain fingerprint

Generating brain fingerprints of subjects, based on their multi-modal data, involves multiple steps: extracting local descriptors in images to build a bag of features (BoF) representation of subjects, building a subject proximity graph by comparing their BoF representations, and embedding this graph in a low-dimensional manifold. Additionally, once the manifold has been constructed, an out-of-sample extension strategy is required to compute the fingerprint of new subjects.

#### 2.2.1. Bag of feature (BoF) representation of subjects

In the first step, a set of local descriptors [40] is obtained from each available image (3D scan). Various local feature extraction and representation approaches [59] can be used, for example, Scale Invariant Feature Transfrom (SIFT) [39] or Speeded UP Robust Features (SURF) [60]. In this work, we use 3D SIFT descriptors as they have been well studied for neuro-image analysis [23, 44, 61] and can be computed efficiently.

3D keypoints are located in the scans of each subject by finding the local extrema (i.e., maxima or minima) of the difference of Gaussians (DoG) occurring at multiple scales. Keypoints with a low contrast or corresponding to edge response are discarded, and remaining ones are encoded into a feature vector (i.e, the descriptor) using the histogram of oriented gradients (HOG) within a small neighborhood. Note that these descriptors are robust to changes in illumination, scale and rotation, and are thus efficient for comparing images acquired using different scanners or imaging parameters. Each subject is then represented as an orderless bag of features (BoF), containing all the descriptors found in this subject’s scans. This representation provides a simple, robust and extensible way of incorporating data from multiple modalities.

#### 2.2.2. Subject proximity graph

Because the BoFs of two subjects may contain different numbers of descriptors, they cannot be directly compared. To circumvent this problem, we construct an intrinsic manifold of subject appearance using a feature-to-feature nearest-neighbor (NN) graph. In this graph, each descriptor is represented by a node and is connected to its K most similar descriptors based on Euclidean distance. This feature-to-feature graph is then converted to a subject-to-subject (i.e., subject proximity) graph by considering, for each pair of subjects, the number descriptors in their BoF that are linked in the feature-to-feature graph.

Let 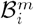 and 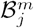 be the BoFs (i.e., set of descriptors) of subjects *i* and *j* for modality 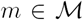, where 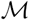 is the set of available modalities. The similarity between these subjects is evaluated as

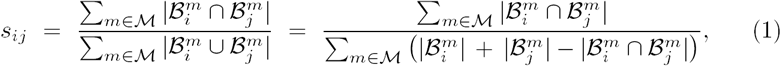

where 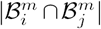 is the number of edges in the feature-to-feature graph between nodes in 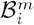 and those in 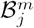. When using a single modality, this measure corresponds to the well-known Jaccard similarity. Here, we extend it to a multimodal setting by comparing the descriptors of each modality separately. We note that the Jaccard distance, defined as one minus the Jaccard similarity, is a *metric* and thus well-suited for constructing the manifold.

When defining the feature-to-feature graph, K determines the number of nearest-neighbor connections for each descriptor. In manifold learning approaches, this parameters controls the locality of the manifold approximation at each point [31]. Its value should be large enough to capture the manifold’s local structure, but also restricted so that distances to nearest-neighbors are close to the geodesic. In our experiments, we tested 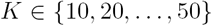 and found similar results for these values. In what follows, we report results obtained with K = 20.

#### 2.2.3. Manifold embedding

A manifold embedding technique is used to obtain compact brain fingerprints from the subject proximity graph. While various approaches could be employed for this task, for instance Isomap [32], locally linear embedding (LLE) [33] and multi-dimensional scaling (MDS) [35], we performed the embedding using Lapla-285 cian eigenmaps [34]. This technique, which is connected to the well-known Laplace-Beltrami operator, has the advantage of being efficient and allowing out-of-sample extensions.

In Laplacian eigenmaps, each subject *i* is mapped to a coordinate vector 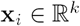 of the manifold, whose dimension *k* is a user parameter. The embedding of subjects in the manifold is made such that two subjects *i* and *j* with a high similarity S*ij* will be close to one another. Let 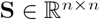 be the adjacency matrix of the subject proximity graph, as defined in Eq (1), and denote as **L = D – S** the Laplacian of **S**, where **D** is a diagonal matrix containing the row sums of **S**. The embedding is accomplished by solving the following problem:

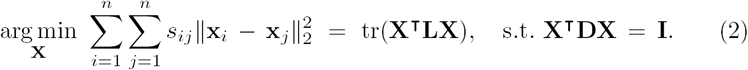

The constraint on **X** removes arbitrary scaling factor in the embedding. As described in [34], the solution to this problem is given by the leading *k* eigen-vectors of the normalized adjacency matrix 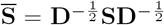, starting from the second one^3^. Once computed, the rows of matrix **X** correspond to the *n* subject fingerprints of size *k*.

#### 2.2.4. Out-of-sample extension

The manifold embedding technique described above computes the fingerprint of all subjects at once. If new subjects are added, this process must be repeated over again, which is inefficient and changes the fingerprint of previous subjects. To alleviate these problems, we use an out-of-sample extension of Laplacian eigenmaps, based on the Nystrom method [62, 63].

Suppose we want to compute the manifold embedding of *m* new subjects. The first step is to update the nearest-neighbor feature graph with the local descriptors of these new subjects, leaving unchanged the nearest-neighbors of the *n* base subjects. We then evaluate the pairwise similarities between new subjects and the base ones. Let 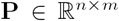 be the matrix containing these similarities, the adjacency matrix of the extended subject proximity graph 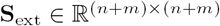 is given by

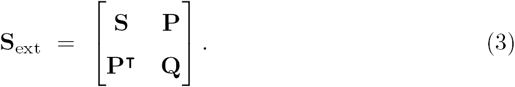

Using the formula in [34], the matrix **Q** of similarities between new subjects can be approximated as **P^T^S^-1^P**.

To normalize **S**_ext_, we compute the vector of row sums

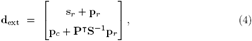

where **s**_r_, 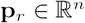 contain the row sums of **S** and **P**, respectively, and 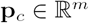 contains the column sum of **P**. In the case where *m* is small compared to *n*, we have that 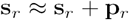, and thus **d**_ext_ can be approximated as

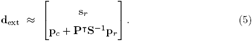

This strategy, used in [64] for white matter fiber segmentation, allows preserving the previous embedding of base subjects. Let **D**_ext_ be the diagonal matrix with entries corresponding to **d**_ext_, the normalized adjacency matrix of the extended subject graph is calculated as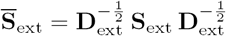. The extended embedding is then obtained following Nystrom’s method as

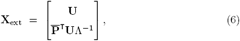

where 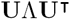 is the eigendecomposition of 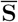, and 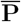 is the normalized submatrix in 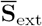 corresponding to **P**. Hence, the embedding of base subjects is the same as in Section 2.2.3, and new subjects are embedded as 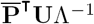. Once more, a fingerprint of size *k* is obtained by considering only the *k* leading eigenvectors in matrix **U**, ignoring the constant eigenvector.

### 2.3 Computational efficiency

Computational and memory requirements are key factors when performing large scale analyses. In this section, we evaluate these requirements for the main steps of the proposed framework. To highlight the efficiency of encoding images with local descriptors, we also compare our framework to a simple fingerprint using full images as features. Other aspects like scan resolution and image alignment requirements are discussed in Section 3. All experiments were performed on a 3.6 GHz processor with 32 GB RAM.

For the BoF representation of images, we extracted 3D SIFT features using a publicly available tool^4^. Computing these features took about 3 seconds per image, and approximately 60 minutes for all 945 images, when processed sequentially. This runtime could however be reduced significantly by processing images in parallel. The feature matching routine [42], for generating the subject proximity graph from the BoF representations of all images, required around 5 minutes to complete. In comparison, calculating the sum of squared distances (SSD) between full images took 1.7 seconds on average for a single pair, and 760,000 seconds for all (945 × 944)/2 = 446,040 pairs (with parallel computations). In terms of memory, each BoF file is approximately 400 KB in size, compared to 84 MB on average for a NIfTI volume file. In summary, the proposed framework is highly efficient in terms of computational and memory requirements compared to a baseline fingerprint using full images. Moreover, since computing the subject proximity graph has a complexity in *O(N* log *N)* where *N* is the number of images, and because extending the manifold embedding can be done efficiently using the Nystrom method, our framework is scalable to large datasets.

### 2.4 Evaluation measures

To measure the link between fingerprint similarity and genetic proximity, we performed a rank retrieval analysis using the sibling information provided in the HCP dataset. In this analysis, we try to identify the twins/siblings of a given subject by comparing its fingerprint with that of all other subjects in the group. Another goal of this analysis is to provide a common platform for the quantitative comparison of individual modalities and their combination. Two standard performance metrics for rank retrieval are used to evaluate the fingerprints: mean recall@k and mean average precision (MAP) [65].

Given a subject *i*, we rank all other subjects by the similarity (i.e., inverse of Euclidean distance) of their fingerprint to that of subject *i*. Denote as 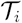the set of target siblings/twins of subject *i*. For instance, if the target group is non-twin siblings (NT), then 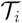 contains the siblings of subject **i** that are not his/her twin. Moreover, let 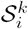 be the set containing the *k* subjects with fingerprints most similar to that of *i* (i.e., the *k* nearest neighbors of *i*). For a given value of *k*, we evaluate the retrieval performance using the measures of recall@k and precision@k:

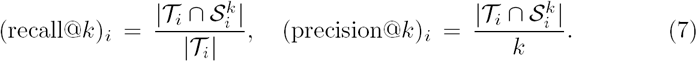

When analyzing the rank performance for a particular sibling type (i.e., monozygotic twin, dizygotic twin or non-twin sibling), we average values over the set of subjects which have at least one sibling of this type, i.e. the set 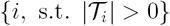.

We also evaluate performance with the average precision, which extends the above metrics by considering the rank of nearest neighbors:

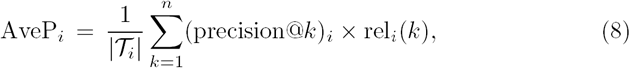

where rel*_i_(k)* is an indicator function with value equal to 1 if the *k*-th nearest neighbor of *i* is relevant (i.e., is in 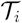), and zero otherwise. The MAP is obtained by averaging AveP values over all subjects having at least one sibling of the target type.

Finally, we use the d-prime sensitivity index [66] to obtain a quantitative measure of separability between the distribution of fingerprint distances corresponding to different sibling types. Let *μ*_1_,*μ*_2_ and *σ*_1_,*σ*_2_ be the means and standard deviations of compared distance distributions (e.g., distance between monozygotic twins versus between dizygotic twins). The d-prime index is computed as

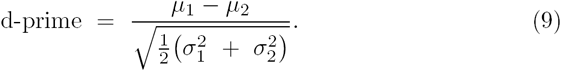

In our experiments, we report absolute values of d-prime, higher values indicating better separability.

## 3. Experiments and results

A comprehensive set of experiments was conducted to analyze the proposed fingerprint and evaluate its usefulness in various applications. In the first experiment, we analyze the manifold embedding of subjects and measure the impact of manifold dimensionality on the fingerprint’s ability to capture genetic proximity. We then perform a detailed rank retrieval analysis, in which fingerprints obtained from a single modality or combinations of multiple modalities are used to identify three types of genetically-related subject: monozygotic twins (MZ), dizygotic twins (DZ) and full siblings (FS). The driving hypothesis of this experiment is that individual modalities capture distinct properties of brain tissues, which can be effectively encoded in the fingerprint, and that combining different modalities can help describe the uniqueness of individual brains. Another goal of this experiment is to measure the relationship between the similarity of fingerprints, for different modality combinations, and genetic proximity.

In another experiment, we assess the impact of following factors on the proposed fingerprint: image alignment, scan resolution, inclusion of skull, and subject age. This is followed by a reproducibility analysis, performed with the restest scans of 42 subjects, and a comparison with a baseline fingerprint using full images as features. The objective of these experiments is to demonstrate the robustness and performance of the proposed fingerprint, compared to a full image scan-based fingerprint.

We also present applications of the proposed framework for identifying retest scans, duplicate corrupt scans, and incorrectly-reported zygosity labels. In addition, we use the segmentation masks provided with the HCP data to identify cortical and subcortical brain regions where the distribution of feature correspondences between monozygotic twins is significantly different from dizygotic twins. In this analysis, we want to find brain regions which are more influenced by genetic promixity. Finally, we conduct a hemisphere asymmetry analysis using the feature correspondences for different types of siblings.

### 3.1. Manifold approximation analysis

To analyze the manifold approximation, we generated fingerprints by projecting the subject proximity graph onto a varying number of spectral components (i.e., leading eigenvectors of the normalized adjacency or Laplacian matrix). Fingerprints were normalized by converting each fingerprint to z-scores (centered to have mean 0 and scaled to have standard deviation 1). Figure 2 (top row) shows a representative 2D spectral embedding of subject proximity graphs obtained using T1w, FA, or both modalities (T1w+FA). As described in Section 2.2.2, modalities are combined by aggregating the feature correspondences in each modality when computing the pairwise subject similarities. In these plots, the position of each red dot corresponds to the 2D fingerprint of a subject. Additionally, in each plot, a single pair of MZ twins is highlighted using blue and cyan dots and their NT sibling highlighted using a green dot.

**Figure 2:**
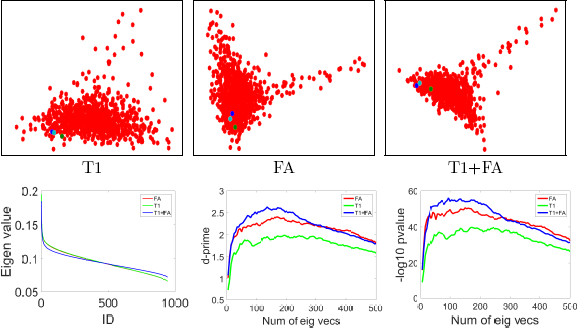
Compact fingerprint analysis. Top row: representative 2D spectral embedding visualization, blue and cyan dots show one pair of MZ twins and green dot shows their not twin (full) sibling; Bottom row: plots of eigenvalues (excluding the first), absolute d-prime, and -log_10_ (p-value) (unpaired t-test) for Euclidean distances between MZ pair vs DZ pair fingerprints with increasing number of eigenvectors.

It can be seen that the distribution of subject embeddings in the manifold varies from T1w to FA, showing that these modalities encode different properties in the fingerprint. Differences between the distributions of FA and T1w+FA fingerprints are in part explained by the fact that spectral embeddings are defined up to a rotation or axis flipping. Moreover, we observe for all three modality combinations that genetically-related subjects are near to each other in the manifold, and that MZ twins are closer than their non-twin (full) sibling.

In Figure 2 (bottom row), we measure the impact of manifold dimensionality on the fingerprint obtained with T1w, FA or T1w+FA modalities. The left plot shows the eigenvalues (sorted by decreasing magnitude) of the subject proximity graph’s normalized adjacency matrix, which reflect the amount of connectivity information captured by their corresponding eigenvector. This plot indicates that most information is encoded in the first leading eigenvectors and, thus, that a compact fingerprint is possible.

This hypothesis is further confirmed in the middle and right plots of the same row, which evaluate for an increasing number of spectral components (i.e., fingerprint size) how the distributions of distances between MZ fingerprints and between DZ fingerprints differ. The separability between these two distributions of fingerprint distances is measured in terms of d-prime (middle plot) and unpaired t-test p-values (in -log_10_ scale). In both measures, higher values correspond to a greater separability. For all three modality combinations, a peak separability is observed around 150 eigenvectors, suggesting this value to be suitable for the fingerprint size. The decrease in separability for larger manifold dimensions is due to the fact that the added eigenvectors encode small variations of brain geometry which are not related to genetic proximity. Nevertheless, the difference between fingerprint distances in MZ pairs and in DZ pairs is significant with p-value < 0.01, for all tested manifold sizes and modality combinations.

Comparing the three modality combinations, the diffusion-based fingerprint using FA images provides a higher separability than the fingerprint generated from T1w, for all manifold sizes. However, the separability is increased further when combining both modalities in the fingerprint, in line with our hypothesisxthat multi-modal fingerprints are more discriminative than those based on a single modality. The impact of modalities on the fingerprint is analyzed in 425 greater details.

Finally, Figure 3 gives the count histograms and probability density curves (fitted) of Euclidean distances between fingerprints of different sibling types. To generate these results, and in all following experiments, we used a fingerprint of 150 features (i.e., leading eigenvectors of the normalized adjacency matrix). Each fingerprint converted to z-scores (centered to have mean 0 and scaled to have standard deviation 1). It can be seen that the fingerprints of MZ twins, which share the most genetic material, are significantly more similar than those of DZ twins or full siblings (FS). This follows the results of various twin studies [50, 13], highlighting the relationship between genetic proximity and anatomical similarity.

**Figure 3:**
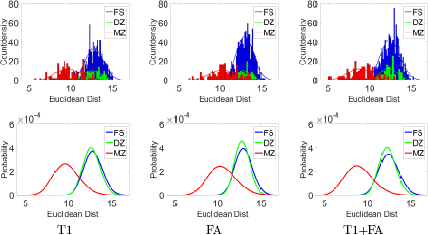
Compact fingerprint comparison for genetically related subjects. Count-density histograms (top row) and probability-normalized curves (bottom row; gamma histogram fitting) of Euclidean distances between twin/sibling pair fingerprints using 100 eigenvectors.

### 3.2 Identification of genetically-related subjects

In this section, we use genetically verified labels of the HCP dataset to determine whether fingerprints generated using different modality combinations can identify genetically-related individuals within a group of subjects. For combining structural and diffusion modalities, we considered data at 1.25mm resolution. For resting state fMRI, we utilize the connectivity matrices (netmats) as functional connectivity fingerprints, and obtain the subject proximity graph (manifold approximation) by computing pairwise Pearson correlation. The idea is to closely follow the functional connectivity fingerprint and similarity computation described in Finn et al. [9] (the parcellation and dataset sizes are different). The multi-modal combinations involving rfMRI are obtained by linear combination of rfMRI subject proximity graph with other modality/combination based subject proximity graph. The weights for linear combination were determined by grid search, and optimal values of evaluation measures are reported. For FreeSurfer based measures, we used the unrestricted csv file, considering volume of sub-cortical structures, thickness and area measures for cortical regions. Each of the measures were converted to zscore across subjects, and then used as a fingerprint (volume measures are first divided by FS-IntraCranial-Vol). Subject proximity graph is approximated by computing the pairwise Pearson correlation. We refer the reader to Section 2.4 for details on the evaluation protocol and measures.

Table 2^5^ reports the mean average precision (MAP) values obtained in a rank retrieval of three different siblings types (MZ, DZ and FS), using fingerprints generated from various modality combinations. Mean recall@k results are reported in supplement material (Figure 1 and Table 8). A rich and diverse set of observations can be drawn from this table. In the next section, it is used to analyze the impact of different factors on the proposed fingerprint’s ability to identify genetically-related siblings, such as scan resolution, image alignment and skull inclusion. While Table 2 includes FA, MD, GFA, rfMRI netmat (partial correlation, ICA 100), and FreeSurfer Volume+Thickness+Area (V+T+A) based mean average precision values, detailed results on dMRI based measures (DTI and GQI), rfMRI netmats, and FreeSurfer measures are described in Table 3, 4, and 5 of supplement material, respectively. Moreover, results on the significance between the distributions of MAP values obtained for different modality combinations and sibling types are also reported in Table 1 of Supplement material.

**Table 2:** Mean average precision (MAP) table comparing different modalities for the task of genetically related subject identification.

| Experiment | Modality | Mean Avg Prec |  |  |
| --- | --- | --- | --- | --- |
|  |  | MZ | DZ | FS |
| sMRI | T1w | 0.886 | 0.160 | 0.128 |
|  | T2w | 0.908 | 0.212 | 0.111 |
| dMRI | FA | 0.964 | 0.219 | 0.160 |
|  | MD | 0.803 | 0.114 | 0.086 |
|  | GFA | 0.968 | 0.234 | 0.161 |
| rfMRI | netmat | 0.968 | 0.352 | 0.205 |
| Modality<br>Combination | T1w+T2w | 0.970 | 0.283 | 0.183 |
|  | T1w+FA | 0.977 | 0.279 | 0.210 |
|  | FA+MD | 0.978 | 0.259 | 0.198 |
|  | T1w+rfMRI | 0.990 | 0.460 | 0.279 |
|  | FA+rfMRI | 0.996 | 0.472 | 0.301 |
|  | T1w+T2w+DTI | 0.994 | 0.392 | 0.270 |
|  | T1w+T2w+FA+rfMRI | **0.997** | **0.546** | **0.371** |
| Skull Impact | T1w Skull | 0.990 | 0.305 | 0.230 |
|  | T2w Skull | 0.980 | 0.310 | 0.164 |
| Alignment Impact | T1w MNI | 0.852 | 0.087 | 0.101 |
|  | T2w MNI | 0.827 | 0.147 | 0.111 |
| Resolution Impact | T1w 1.25mm | 0.831 | 0.136 | 0.121 |
|  | T2w 1.25mm | 0.879 | 0.173 | 0.132 |
| Baseline Comparison | T1w | 0.649 | 0.079 | 0.052 |
|  | T2w | 0.520 | 0.069 | 0.038 |
|  | FA | 0.707 | 0.076 | 0.049 |
|  | V+T+A (FreeSurfer) | 0.795 | 0.172 | 0.106 |
| Retest set | T1w | 0.915 | 0.137 | 0.130 |
|  | T2w | 0.917 | 0.212 | 0.113 |
|  | FA | 0.944 | 0.252 | 0.158 |
| Random | Rand | 0.005 | 0.005 | 0.006 |

**Table 3:**
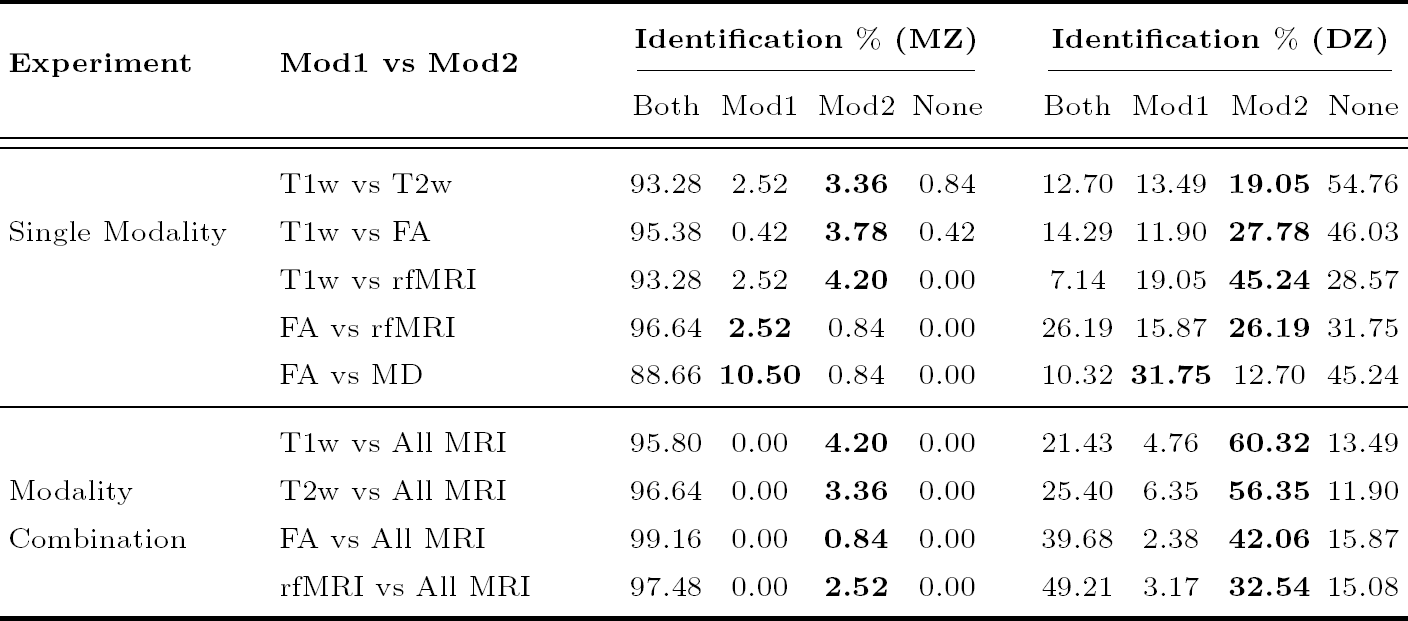
Relative informativeness of fingerprints from two modalities. Comparison between modalities or their combination for the task of identification of a given sibling type. The reported values are **relative percentages** of MZ/DZ twin identification for two modalities, with Mod1 representing successful identifications by modality 1 only. The total number of identification tasks is 238 and 126 for MZ and DZ respectively. Note: identification of twin 1 by twin 2 and vice-versa are considered two separate tasks. The identification is considered a success if the twin is identified within the 10 nearest neighbors of a subject (among 944 subjects). (All MRI= T1w+T2w+FA+rfMRI)

**Table 4:**
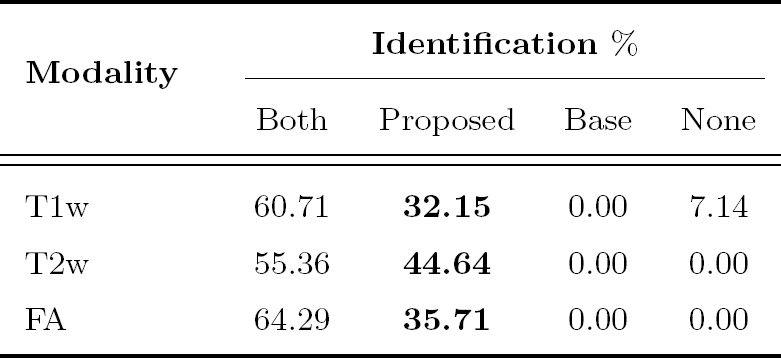
Analysis of self-reported zygosity to genetically verified zygosity detection. Relative percentage of DZ-to-MZ subject’s twin identification within 10 nearest neighbors for proposed framework vs Baseline (Full image based pairwise SSD). Total number of identification tasks is 56. Identification is considered a success if the twin is identified within the 10 nearest neighbors of a subject.

**Table 5:** Significant parcellations for T1w and T2w for MZ vs DZ and MZ v FS, along with the HolmBonferroni corrected p-values (-log_10_ scale) obtained using unpaired t-test (Feature match count). **corrected p-values < 0.05** are in bold.

| Label | T1w |  | T2w |  |
| --- | --- | --- | --- | --- |
|  | MZ vs DZ | MZ vs FS | MZ vs DZ | MZ vs FS |
| subcortical | **29.31** | **50.31** | **26.06** | **39.41** |
| Crtx-LH | **22.85** | **35.17** | **23.37** | **38.87** |
| Crtx-RH | **21.48** | **39.76** | **25.38** | **37.73** |
| WM-LH | **37.64** | **62.83** | **27.88** | **47.38** |
| WM-RH | **23.38** | **36.80** | **21.88** | **32.62** |
| L-Lat-Vent | **5.84** | **11.34** | **5.31** | **7.50** |
| R-Lat-Vent | **4.21** | **10.72** | **3.99** | **7.57** |
| L-VentralDC | **1.49** | **6.06** | **5.16** | **2.98** |
| R-VentralDC | 0.45 | 0.60 | 0.00 | 0.01 |
| R-Cerebellum-WM | **3.98** | **15.40** | 0.00 | 0.57 |
| L-Cerebellum-WM | **4.82** | **11.11** | **2.33** | **6.55** |
| R-Putamen | 0.48 | 0.51 | **2.34** | 1.24 |
| L-Putamen | 0.87 | 0.35 | 0.06 | 0.30 |
| L-Cerebellum-Crtx | **5.74** | **6.26** | **5.58** | **13.81** |
| L-Thalamus-Proper | **2.71** | **4.03** | 0.37 | 0.01 |
| 4th-Ventricle | **1.49** | **2.24** | **1.86** | **3.91** |
| L-Hippocampus | **3.23** | **3.76** | **4.61** | **5.85** |
| CC-Anterior | **1.83** | 0.51 | 0.40 | 0.71 |
| R-Cerebellum-Crtx | **5.96** | **11.93** | **3.38** | **7.57** |
| 3rd-Ventricle | 0.45 | 0.51 | 0.37 | 0.43 |

Comparing modalities, we observe that rfMRI netmat yields the highest MAP among all single-modality fingerprints, with a significant margin for DZ and FS. For structure-based fingerprints, T1w and T2w provide similar performances across the different sibling types, with MZ and DZ MAP values being higher for T2w. Similarly, for diffusion based fingerprints, FA and GFA provide similar performance, while outperforming MD. Furthermore, higher MAP values are obtained when combining multiple modalities, the combination of T1w, T2w, FA, and rfMRI having the best performance for all sibling types. This applies for combinations within/across structural or diffusion modalities: T1w+T2w outperforms T1w and T2w, FA+MD performs better than FA and MD, T1w+FA outperforms T1w and FA, etc. Similarly, T1w+rfMRI outperforms T1w and rfMRI, and FA+rfMRI performs better than FA and rfMRI.

With respect to the tested sibling types, we observe a mean average precision between 80.3% and 99.7% when using the fingerprint to identify MZ twins, for all modalities and their combinations. This illustrates the high impact of genetic similarity on both structural and diffusion geometry in the brain as well as functional connectivity. Comparing MZ, DZ and FS siblings, we see higher MAP values for MZ twins compared to DZ twins or full siblings, supporting the fact that MZ twins share more genetic information [67]. In contrast, performances obtained for DZ twins and full siblings are comparable, which reflects the fact that both sibling types have the same genetic proximity. In general, the differences between DZ twins and full siblings were found to be not significant in unpaired t-test for individual modalities, with T2w being the exception (Supplement material Table 1). Also, similar observations can be drawn from mean recall@k plots and mean recall@10 values (supplement material Figure 1 and Table 8), with added information on difference in modalities with increasing *k*. Mean recall@k, for k = 1,…, 50, also known as sensitivity, evaluates the proportion of individuals that are genetically related to a given subject, which are within the k individuals most similar to that subject (in terms of fingerprint distance). In addition, we observe higher MAP values for full sibling identification vs maternal half sibling (MHS) identification (supplement material Table 6). While MAP values for paternal half sibling identification show lot of variation, mainly due to limited sample size.

To quantify the informativeness of one modality versus another, Table 3 reports the relative percentage of MZ and DZ twins identified by both, a single, or none of the modalities^6^. Note, for a given twin type, each row provides relative comparison between two modalities, with sum of row being 100%. The total number of identification tasks is 238 for MZ and 126 for DZ (the identification of twin 1 by twin 2 and vice-versa are considered two separate tasks). For each task, we consider the *k* = 10 nearest neighbors of a subject in terms of fingerprint distance. The identification is considered a success if the subject’s twin is identified within these neighbors. When comparing the relative success rates of single modalities (top half of the table), we observe that FA identifies more twins uniquely than when using T1w or MD. This is particularly noticeable for DZ twins, where 27.78% of DZ pairs were identified by the FA-based fingerprint but not the T1w-based ones. Yet, structural modalities still capture brain tissue properties that are not provided by dMRI, as shown by the 11.90% of all DZ pairs that are identified using T1w but not with FA. Similar observations can be drawn when comparing rfMRI with structural and diffusion modalities. For example, rfMRI identifies 45.24% of DZ pairs that are not identified using T1w within 10 neighbors, and T1w identifies 19.05%.

As with the results in Table 2, we see that combining multiple modalities leads to a more discriminative fingerprint. For example, 4.20% of MZ and 60.30% of DZ twins are identified by fingerprints generated from all modalities (i.e., All MRI=T1w+T2w+FA+rfMRI) but not from fingerprints based only on T1w. Reversely, all MZ twins identified with T1w are also found using T1w+T2w+FA+rfMRI, and only 4.76% of DZ twins are identified uniquely with T1w. This last result could be explained by the fact that subjects can have local similarities due to factors not related to genetics.

### 3.3. Impact of various factors

Factors like image alignment, scan resolution, skull inclusion and subject age, can be confounds when analyzing the proposed fingerprint. In the following subsections, we measure the impact of these factors on the fingerprint’s ability to find genetically-related subjects.

#### 3.3.1. Image alignment

Population-level analyses usually require aligning images to a common space or segmenting them into regions of interest, two steps which can be computationally expensive.

Table 2 (sMRI vs alignment impact rows) reports the retrieval performance obtained for fingerprints generated from T1w and T2w images in MNI space (0.7mm resolution, data provided by the HCP with affine alignment to MNI template). For all sibling types, MNI space-aligned fingerprints (denoted as MNI in the table) obtained lower MAP values than fingerprints using native space data. This observation, which is consistent across T1w/T2w modalities and all sibling types, indicates that image alignment is not required for our fingerprint. Although the difference is not significant, in general, in an unpaired t-test with p-value < 0.01 (see Table 2 of Supplement material), bringing subjects to a common space may reduce the discriminativeness of the fingerprint leading to the reduction in MAP values. Note that similar results were obtained using full images as fingerprints (analyzed in the following section), with lower MAP for affine-aligned images.

#### 3.3.2. Scan resolution

Scan resolution is another important factor in multi-modal and multi-subject analyses, for example, sMRI data usually offer higher resolutions compared to dMRI.

Table 2 (sMRI vs resolution impact rows) shows that MAP values for the MZ/DZ twin identification task decrease when going from 0.7mm to 1.25mm resolution, for both T1w- and T2w-based fingerprints. This is due in part to the reduced number of SIFT features extracted from 1.25mm resolution image, compared to 0.7mm resolution ones. However, this is not the case for FS identification tasks, where contrasting trends are observer for T1w and T2w. Moreover, differences in MAP values for the two resolutions are not significant when running an unpaired t-test with p-value < 0. 01, for any sibling type (see Supplement material). These results suggest the robustness of our framework to varying scan resolutions.

#### 3.3.3. Inclusion of skull

Since skull size and shape is strongly influenced by genetics, including skull information in fingerprints could increase their discriminative power. In this experiment, we measure the usefulness of skull tissues for identifying pairs of MZ, DZ and FS subjects. (Facial features are not analyzed)

Table 2 reports the performances of fingerprints based on T1w and T2w image, with or without skull stripping. For both T1w and T2w, as well as all sibling types, including the skull in images improves MAP values. These results are significant, with p-value < 0.01, in an unpaired t-test (see Table 2 of Supplement material). Hence, skull tissues provides additional feature correspondences which help identify twins and non-twin siblings. However, we should mention that skull stripping is essential to most neuroimaging analyses, and our objective here is only to measure the informativeness of skull tissues on the proposed fingerprint. An extended skull-inclusion analysis, including T1wbyT2w MRI ratio images (myelin content) and modality combinations are reported in supplement material, Table 10.

#### 3.3.4. Subject age

In twin studies, the age of subjects can be a confound when comparing between different sibling types. For instance, DZ twins and FS siblings share the same amount of genetic material, yet DZ twins could be more similar due to their same age. The HCP data used in this study was acquired in the age range of 22-36, which corresponds to the plateau/saturation in brain and white matter development [52, 51]. Nevertheless, we analyze whether age differences in non-twin siblings is a contributing factor on performance.

Toward this goal, we divided FS sibling pairs in two groups based on the median age difference of 3 years, and measured the MAP in each group, for fingerprints generated from T1w, T2w, and FA. Similarly, we also evaluated the impact of absolute age on performance of MZ/DZ. In this case, we divided subjects (not subject pairs) in two groups based on the median subject age of 29 years. As shown in supplement material Table 9, no significant differences in MAP are observed across these groups. In summary, using the HCP dataset, we found no significant impact of subject age on the proposed fingerprint.

### 3.4. Comparison to baseline fingerprint

We compared the performance of our fingerprint to a baseline using full images as features. In this baseline, the similarity of two fingerprints is measured as the sum of squared distances (SSD) between intensities of corresponding voxels. Table 2 gives the MAP obtained using this baseline, for T1w, T2w, and FA images in native subject space. For MZ twin identification, the baseline using FA images performs better than T1w or T2w images, which is consistent with the results of the proposed fingerprint. However, we see that our fingerprint performs consistently better than the baseline, with MAP improvements of 0.237 in T1w, 0.388 in T2w, and 0.257 in FA, for the task of identifying MZ twins. These improvements are significant in a one-sided unpaired t-test with p-value < 0.01 (see Supplement material, Table 2). Note that we also tested a similar baseline created from MNI aligned images, however this led to lower MAP values.

In addition, we use Freesurfer derived measures of sub-cortical volumes, and thickness and area of cortical regions as baseline fingerprints (see Supplement material Table 5 for detailed analysis on FreeSurfer measures). Higher MAP values are obtained for MZ twin identification using our fingerprint vs Vol+Thck+Area FreeSurfer (0.886 vs 0.795, p-value < 0.01). However, no significant difference is observed for DZ and FS identification.

In summary, while being very compact and efficient (see Section 2.3), our fingerprint based on local features is significantly more informative than a voxel-based representation and performs similar/better than FreeSurfer derived measures for identifying genetically-related subjects.

### 3.5. Results reproducibility

To test the reproducibility of the results, we re-ran the same analysis after replacing the T1w, T2w and FA images of 42 subjects with their retest data. Table 2 gives the MAP values obtained following this process. We observe small differences in MAP, compared to fingerprints using the original data, however, these are not significant (see Supplement material Table 2).

We note that the majority of retest subjects available in the HCP data are MZ twins. Since we do not observe significant differences for identifying this type of twins, it shows that the results are reproducible. The small differences in MAP values for DZ twins and FS siblings could be attributed to slight changes in the ordering of retest subjects’ nearest neighbors.

### 3.6. Applications

In this section, we demonstrate the usefulness of our fingerprint on three different applications: 1) the correction of erroneous zygosity labels, 2) the detection of retest and duplicate scans, 3) the visualization and analysis of local feature correspondences for different modalities, sibling types and neuroanatomical regions.

#### 3.6.1. Zygosity label correction

The Q3 release of the HCP dataset contained self-reported zygosity labels for twins. In the HCP 1200 release, which contains genetically verified zyosity labels, it was found that many self-reported DZ twins were actually MZ twins. In light of this problem, we first evaluate if the proposed framework can be used to identify the twins in large dataset whose self-reported zygosity differs from their true zygosity.

In earlier experiments, we found higher MAP values for MZ and that the MZ twin of subjects was within the 10 nearest neighbors of a subject (i.e., a mean recall@k of 100% was obtained for k ≤ 10, supplement material Table 8), regardless of the modality combination used for the fingerprint. Conversely, a lower percentage of DZ twins could be identified in these lists of nearest neighbors. Based on this idea, we find incorrectly reported MZ candidates as the DZ twins which are within the 10 nearest neighbors of a subject.

Table 4 reports the percentage of DZ-to-MZ twins (56 in total) correctly identified by the proposed fingerprint, the baseline using full images, both or none of these methods, for T1w, T2w and FA modalities. The results show that our fingerprint can identify most incorrectly self-reported MZ twins, with a detection rate of 92.86% for T1w, 100.00% for T2w, and 100.00% for FA. For all modalities, over 32% of cases were identified uniquely by our fingerprint. In contrast, no DZ-to-MZ twins were identified uniquely by the baseline fingerprint. In conclusion, the proposed fingerprint can be used effectively to detect misreported zygosity labels.

#### 3.6.2. Retest and duplicate scan identification

To analyze our fingerprint’s ability to detect repeat scans of the same subjects (acquired after a time gap), we used the data of 945 subjects + 42 retest subjects, and considered the task of identifying repeat scan in a rank retrieval analysis.

Following the same evaluation protocol as for identifying MZ/DZ/FS siblings, we obtained a MAP value of 1 for fingerprints generated from T1w, T2w or FA. Thus, in all cases, the single most similar fingerprint to that of a subject corresponded to this subject’s retest data. Moreover, when considering the number of local feature correspondences in the subject similarity (i.e., 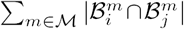 in Eq (1)), we observed more correspondences for the retest data of a subject than for the subject’s MZ twin.

Duplicate scans in a dataset, for example resulting from noise corruption, renaming or other manual errors, can introduce bias in analyses. Therefore, we also assessed if our fingerprint could detect duplicate scans of the same subject, corrupted by noise. For this experiment, we introduced duplicate scans for 42 T1w images, to which was added random noise (uniformly distributed random numbers in the [-a, a] range, where 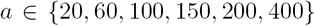 the mean and stdev of image intensities are respectively 720 and 185).

Running a rank retrieval analysis using duplicate scans as target, we again obtained an MAP value of 1, for all tested noise levels. As in the retest scan identification task, the number of local feature correspondences was higher with corrupted duplicates than with images of MZ twins. Compared to retest scans, the number of feature correspondences was nearly half for corrupted duplicated, suggesting that noise can reduce correspondences to some extent. Overall, the results of this experiment demonstrate that our fingerprint can preserve brain characteristics over different scans of a subject.

#### 3.6.3. Local feature correspondence analysis

To understand the advantages and limitations of a BoF-based fingerprint compared to voxel-wise or shape-based methods, we perform an in-depth analysis of local feature correspondences between subjects. In order to compare our findings with those of related fingerprint studies like Brainprint [8], we limit our analysis to genetically-related subjects from HCP and to structural MRI modalities. Other applications of BoF representations for neuro-image analysis have been well studied in the literature [23, 44, 61].

We start with a qualitative visualization of pairwise feature correspondences between subjects of different sibling types. The distribution of correspondences in these modalities is then analyzed using the segmentation maps (WM parcellation) files provided with HCP data. Furthermore, we also report cortical and subcortical regions having significantly different match distributions across sibling types, these regions having a closer relationship to genetic proximity. Finally, we perform a lateral asymmetry analysis in which the distribution of correspondences in hemispheres are compared. Since we perform pairwise matching, an extended set of 1010 HCP subjects: 139 MZ pairs, 72 DZ pairs, and 1214 full sibling pairs.

##### Scale-space visualization of features correspondences

Analyzing local feature correspondences between sibling pairs provides information in terms of their location as well as scale. In 3D SIFT features, scale corresponds to the variance of a Gaussian blur kernel for which the corresponding voxel in the blurred image is a local extrema [40, 39]. It thus coincide, to a certain degree, with the size of structures in which these features are located.

Figure 4 gives a scale-space visualization of features matched between a subject and his/her MZ twin, as well as the subject’s non-twin (full) sibling, for T1w, T2w and FA images (See supplement material for DZ and non-twin (full) sibling). The scale information is represented using the circles’ radius. Note that circles represent the intersection of 3D spheres with the visible slice and, thus, non-intersecting features are hidden in this 2D visualization.

**Figure 4:**
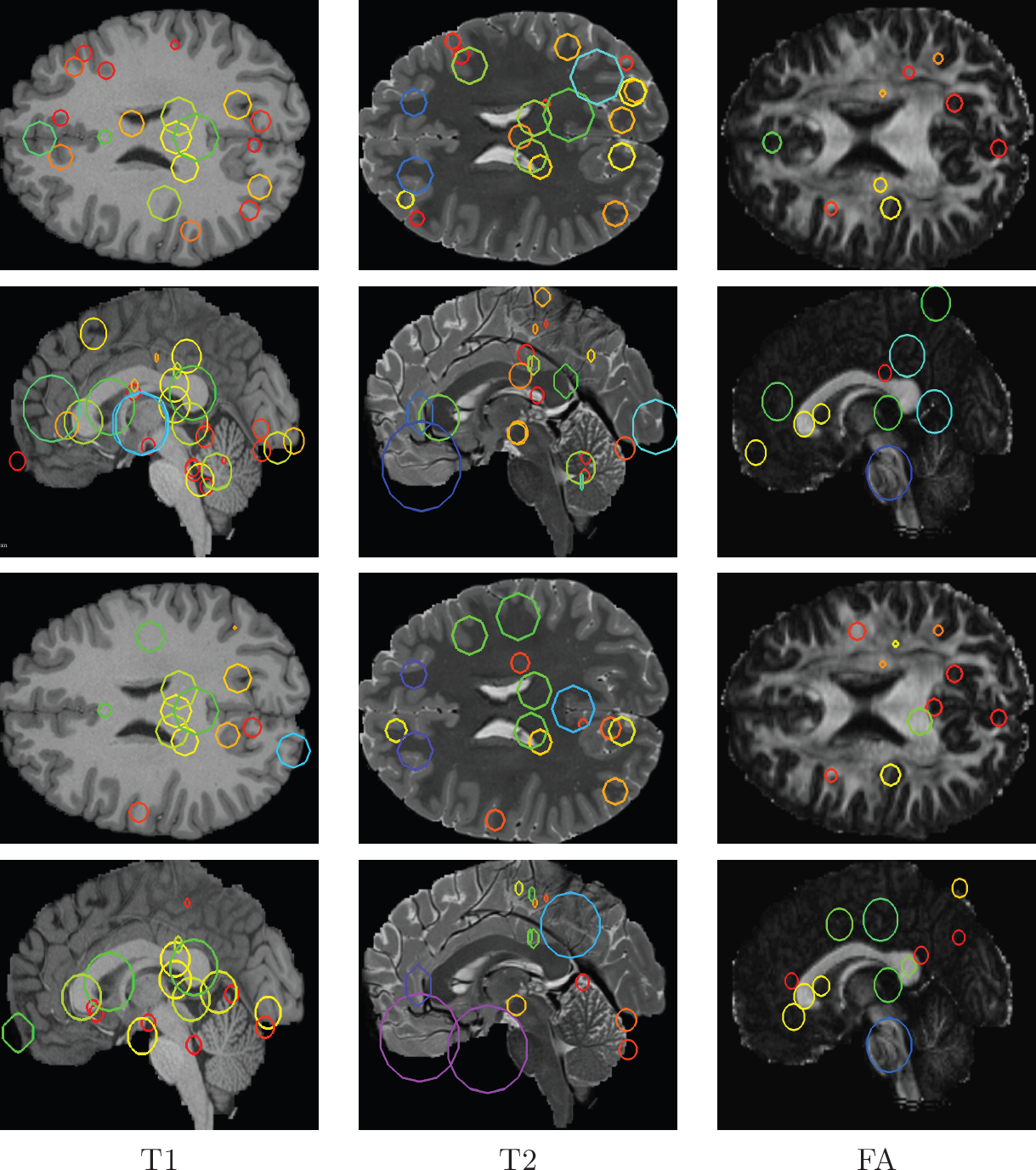
Example of feature correspondences for a subject and his/her MZ twin (rows 1-2), and the subject’s full sibling (FS) (rows 3-4). Scale space is represented using circle radius (for the visible slice).

It can be seen that different image modalities generally result in distinct, complementary feature correspondences throughout the brain. In T1w and T2w images, features are mainly located in the frontal lobe, corpus callosum and cerebellum. Smaller-scale features are also visible along various cortical regions, as well as in subcortical structures near the basal ganglia. Moreover, images based on diffusion measures have less correspondences than in structural modalities. These correspondences are located mostly inside or near to white matter: larger-scale features in the corpus-callosum, and smaller-scale ones in the brain stem and along white matter bundles. The distribution of features in prominent brain regions is further analyzed in the next section.

Comparing different sibling types, we see a greater number of correspondences between MZ twins than between DZ twins or full siblings. This observation, which is easier to visualize in T1w and T2w images, is consistent with other analyses on twin datasets. In terms of feature location and scale, we observe a slightly higher number of correspondences in the frontal cortex for MZ twins, however, no obvious pattern can be drawn from one set of representative plots.

##### Region-wise analysis of feature correspondences

Here, we analyze the distribution of feature correspondences across atlas-defined neuroanatomical regions, measured over the entire group of subjects. For each scan, segmentation labels were obtained from the Freesurfer-processed data, using LUT table for label descriptions.

Figure 5 shows the box plot distributions of feature correspondences between pairs of MZ, DZ and full siblings, and for T1w and T2w images. Feature match counts are reported for five broad regions: non-white matter subcortex (`s-cort`), left/right cortex (`crtx-lh/rh`) and left/right white matter (`wm-lh/rh`). Note that mapping local features to a finer cortical parcellation is difficult due to the limited thickness of the cortex. Subcortical regions are further analyzed below.

**Figure 5:**
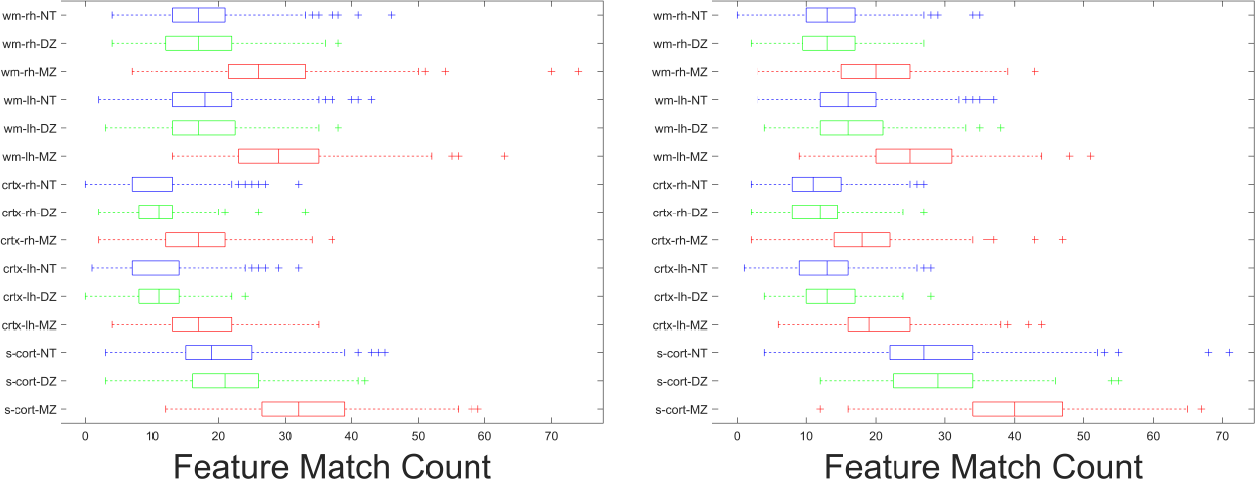
Box plot comparison between MZ, DZ, and FS for pairwise feature correspondence counts for T1w (left) and T2w (right) for major structures. Red, green and blue correspond to MZ, DZ, and FS respectively.

Comparing across sibling types, we observe a higher number of feature correspondences for MZ pairs across all five regions and both T1w and T2w modalities. This confirms once again that the local features employed in our fingerprint captures brain characteristics related to genetic proximity. Analyzing the region-wise distribution of feature correspondences, all five regions are well represented. Since the number of local features in a region is proportional to its size, it is not surprising that the cortex has the least correspondences. Yet, such features are also produced by intensity variations (i.e., edges), thus explaining why many correspondences are found in the cortex. Finally, when comparing T1w and T2w modalities, we see small differences in the match counts, however these are not statistically significant.

To identify regions showing a strong relationship to genetic proximity, Table 5 gives the p-values (-log_10_ scale) of an unpaired t-test comparing the mean number of correspondences between subjects of a given sibling type versus another sibling type (e.g., MZ vs DZ). Significance values are provided for the five major regions described above, as well for 15 prominent subcortical structures matching the analysis by Wachinger et al. [8]. To account for multiple comparisons (i.e., one for each tested region), reported p-values have been corrected using the Holm-Bonferroni procedure [68]. Moreover, to account for age and size bias in this analysis, we selected FS pairs with less than 3 years age difference, and matched the number of FS pairs to MZ pairs using a simple bipartite matching based on age.

From Table 5, we observe significant differences between MZ twins and DZ-twins/full-siblings (-log_10_(p-value) > 2), for all five major regions and for both T1w and T2w images. In subcortical structures of T1w images, cerebellum white matter and cortex (left and right), lateral ventricles (left and right), left hippocampus and left thalamus proper have a significantly different number feature correspondences in MZ twins than in DZ twins or FS subjects. Comparing results obtained with T1w and T2w, the same structures are significant across both modalities, differences in significance reflecting the complimentary of these modalities.

##### Hemisphere asymmetry analysis

In our last experiment, we analyze the symmetry of feature match counts across brain hemispheres, for major structures. Toward this goal, we considered only right-handed (RH) subjects, and limited sibling pairs to subjects with same gender (i.e., a male and his brother, or a female and her sister). For non-twin siblings, we also restricted our analysis to subject pairs with less than 3 years of age difference.

Table 6 gives the results of two-sided unpaired t-tests comparing the feature match counts between cortical or white matter regions (Freesurfer LUT labels) in left- and right-hemispheres. To analyze gender effects, we also report results individually for RH male siblings and RH female siblings. Overall, we observe significant asymmetry in white matter regions (with -log _10_ (p-value) > 2) of MZ twins, the highest significance values obtained for T2w images. No clear pattern is found across sibling types, although hemispherical differences are generally higher in MZ twins than in DZ twins or full siblings. Likewise, no conclusion can be drawn when comparing results for male and female sibling pairs, with significance values varying across different sibling types and modalities.

**Table 6:** Hemisphere asymmetry analysis. For a given modality and twin type we compare feature match count differences across hemisphere for major structures. (unpaired t-test + feature match count)

| Modality | Type | RH Female |  | RH Male |  | RH Pairs |  |
| --- | --- | --- | --- | --- | --- | --- | --- |
|  |  | Crtx | WM | Crtx | WM | Crtx | WM |
| T1w | MZ | 0.95 | 1.20 | 0.15 | **2.57** | 0.87 | **2.73** |
|  | DZ | 1.05 | 0.78 | 0.35 | 0.99 | 1.12 | 0.06 |
|  | FS | **1.71** | 0.39 | 0.84 | 0.11 | **1.89** | 0.09 |
| T2w | MZ | **1.95** | **9.13** | **1.52** | **5.77** | **3.00** | **13.97** |
|  | DZ | 1.06 | **3.60** | 1.29 | 1.23 | **1.93** | **4.06** |
|  | FS | 1.04 | 1.22 | 1.23 | **5.35** | **1.90** | **5.84** |

The asymmetry of function in the brain, for example the hemispheric specializations of language and motor functions, has been extensively studied [69]. Similarly, studies have analyzed anatomical brain asymmetries based on voxel-based morphometry, sulci and other brain features [8].

The multi-modal and multi-region analysis presented in this work extends previous studies of brain asymmetry in the literature by considering sibling types. Accounting for various confounds, including gender, genetics, handedness and age, this analysis has shown a greater asymmetry in feature correspondences between MZ twins than DZ twins and full siblings, mostly found in white matter regions and T2w images. Moreover, differences in asymmetry appear to be directional.

## 4. Discussion

In this section, we summarize the findings of this study and emphasize their link to previous investigations. We also highlight its limitations and discuss additional considerations.

### Identification of genetically-related subjects

Our experiments on the task of identifying genetically-related subjects led to various useful observations. We established that the proposed fingerprint, generated from individual modalities or their combination, respects the genetic relationships between siblings, with MZ twins being more similar than DZ twins or full siblings [50, 13].

3D SIFT detects local points of extrema (minima or maxima) based on changes in the magnitudes in a given image. For example, for T1w the features are located at the boundaries of white matter and grey matter, including subcortical structure boundaries, as shown in feature correspondence visualization. Abstractly speaking it draws discriminative blob like structures while considering scale-space. The intuition is that these discriminative blob like structures capture the key points of change in a given image, a representative summary in scale-space. As such matching them leads to better identification as opposed to voxel-wise full image comparison (say using sum of squared distance) and performs comparably to Freesurfer derived measures of volume, thickness, and area, as well as rfMRI based netmats.

Analyzing the manifold approximation, we also showed that a discriminative fingerprint could be obtained with only 150 spectral components (i.e., leading eigenvectors of the normalized adjacency matrix of the subject proximity graph). When compared to a baseline using full images as features, this compact fingerprint yielded significantly better performances, for all modalities and sibling types. This illustrates the high efficiency of our fingerprint and its advantages for comparing large groups of subjects. Moreover, while Laplacian eigenmaps were used to embed the subject proximity graph, the proposed framework is generic and other approaches (e.g., see [31]) can be employed for this task.

The comparison of fingerprints obtained from structural MRI, diffusion MRI, and resting state fMRI highlighted the informativeness and complementarity of these modalities.

Among individual modalities, resting state fMRI based fingerprint performed best for DZ/FS identification and had similar performance to FA/GFA for MZ twin identification. We hypothesize that good performance of rfMRI netmat is due to the discriminative power of connectivity profiles, which is a result of integration over a relatively long period of time (4800 volumes, and 4 runs of 15 minutes each), as mentioned in Finn et al. [9]. Also, the connectivity profile is based on certain parcellation of brain, adding addition information about individual variability. Moreover, while the MAP values for FA/GFA are similar to rfMRI based MZ twin identification, mean recall@10 and relative identification % showed that FA perfroms slightly better than rfMRI (2.54% unique MZ pair identification as opposed to 0.84% pairs.

We hypothesize that the better performance of BoF based FA fingerprint is due to the framework capturing the changes in contrast/magnitude in FA maps. Since it conveys certain information about white matter bundles and their boundaries in addition to separation between grey matter and white matter, FA performs better than T1w and T2w as well as rfMRI (for MZ twins). The distribution of feature matches in the visualization of one pair of feature correspondences shows this qualitatively.

Furthermore, results of this study demonstrate the usefulness of combining multiple modalities in a brain fingerprint. Thus, better performances were obtained with a combined set of modalities than with these modalities alone. Our results are consistent with previous studies underlining the benefit of a multimodal fusion [24, 25]. As a note, we have focused on major observations only, the comprehensive analysis is open to various other observations including comparison of DTI vs GQI measures, inclusion of T1wByT2w MRI ratio images, FreeSurfer measures based identification, etc.

Finally, the fingerprint proposed in this work is motivated by the recent increase in multi-modal brain studies. Multi-modal MRI has been shown useful for the analysis of neurodegenrative disorders [24]. For example, combining multimodality data also achieves a higher classification performance for identifying subjects with schizophrenia [70].

### Applicability of the proposed fingerprint

Our factor impact analysis demonstrated the robustness of the proposed fingerprint to the non-alignment of images. Furthermore, since image alignment is key for most population level analysis [18], by alleviating this requirement, the proposed fingerprint may help save computational costs and avoids errors introduced during alignment.

Experiments have also shown that scan resolution does not have a significant impact on results, although using lower resolution images reduces the number detected features. Data acquired from multiple sites or scanners often need to be brought to same resolution, introducing small errors during interpolation and re-sampling. The proposed fingerprint may thus be of help for multi-site studies, and pave the way to resolution-independent analyses.

Using retest scans led to no significant changes in results, further validating the robustness of our fingerprint to image acquisition. However, a detailed longitudinal analysis with longer between-scan times would be required to fully confirm this claim.

Similar to the computer vision challenges, the proposed rank retrieval analysis highlight that using twin identification task and MAP as evaluation measure, we can compare brain fingerprints from various modalities and/or their combination. We believe that this could be utilized in future studies. We used the proposed fingerprint to find incorrectly reported zygosity labels and identify retest/duplicate scans of the same subjects. Hence, our fingerprint could serve as efficient and reliable tool for detecting inconsistent information in large cohorts. Another potential application could be to provide physicians with related cases in clinical settings like MCI diagnostic assistance [71].

While various twin studies have analyzed genetic influences based on volume, cortical thickness, surface area, and morphometry [8], this is the first work to use local features and manifold approximation for this problem. Analyzing the distribution of features correspondences across brain regions, in images of different modalities, reveals many interesting insights. Results identify various neuroanatomical regions (e.g., cerebellum, lateral ventricles, ventral diencephalon, hippocampus and thalamus proper) having significantly different match counts in MZ twins than DZ twins or full siblings. These findings relate to those reported in [8], which were obtained on a different dataset (mean subject of age of 56 years, compared to a median of 29 years in the HCP dataset). Another key aspect of our analysis is the size of the subject cohort, larger than that of related studies [50].

### Additional considerations

In this work, we used a rank retrieval analysis to evaluate the relation between fingerprint similarity and genetic proximity. However, estimating heritability directly, for instance using the approach described in [49], would provide a better quantification of genetic influence on fingerprint features. Heritability of multi-modal brain imaging phenotypes, using a more than 8, 000 subjects from UK Bio-bank [17], has been studied in [48]. Similarly, [47] report hearitability of multi-modal functional connectivity using 800 HCP subjects. An extensive analysis will be required to asses heritability of compact fingerprints and relate the findings with these state-of-the-art studies.

Moreover, when building the subject proximity graph, we assumed the independence of feature correspondences across modalities. However, a deeper analysis could be carried out to investigate false feature correspondences and correlation between features correspondences across modality. As mentioned before, other manifold embedding methods like Locally Linear Embedding (LLE) [33] could also be employed for this step.

In this study we analyzed data from sMRI, dMRI and rfMRI. However, the proposed framework is generic and could be extended to other modalities like task-fMRI, PET-MRI and quantitative T1/T2 maps. Finally, this study focused on comparing and combining different modalities for identifying genetically-related subjects, misreported zygosity labels and duplicate/restest scans. An interesting extension of this work would to be to assess whether our fingerprint can be used as a biomarker to identify subjects with cognitive or neurological disorders. Publicly available data, for instance from the ADNI dataset [23] or Parkinson’s Progression Markers Initiative (PPMI) dataset [72], could be used for this analysis.

## 5. Conclusion

We presented a brain fingerprint, based on manifold approximation, for the multi-modal analysis of genetically-related subjects. In a rank retrieval analysis, mean recall@k and mean average precision were used to measure the relation between fingerprint similarity and genetic proximity, as well as the contribution/complementarity of information from different MRI modalities. Results indicated that a compact fingerprint of only 150 features could identify genetically-related subjects better than a baseline using full images as features. Our experiments also showed that each modality provides complementary information which can uniquely identify some sibling pairs. Furthermore, we demonstrated the benefit of considering multiple modalities in the fingerprint, combined modalities leading to a better performance than considering these modalities separately. Moreover, our analysis demonstrated the robustness of the proposed fingerprint to various factors, including image alignment, scan resolution and subject age. The reproducibility of results was also confirmed using retest scans from the HCP dataset, showing our fingerprint to be robust to variability in image acquisition.

The usefulness of our fingerprint was assessed on the tasks of identifying incorrectly reported zygosity and retest/duplicate scans in large dataset. Results of this experiment highlighted the effectiveness of our fingerprint, with MAP values near 100% for all test cases. Moreover, analyzing the distribution of features correspondences across the brain revealed neuroanatomical regions (e.g., cerebellum, lateral ventricles, ventral diencephalon, hippocampus and thalamus proper) with significantly different match counts in MZ twins compared to DZ twins or full siblings. This work could be extended by further investigating the differences, in terms of feature location and similarity, between dizygotic twins and non-twin siblings. A deeper analysis of aging effects could also be performed, for instance, using longitudinal data. Such analysis would help understand the effect of neuroplasticity on individual brain characteristics.

## Acknowledgements

Data were provided in part by the Human Connectome Project, WU-Minn Consortium (Principal Investigators: David Van Essen and Kamil Ugurbil; 1U54MH091657) funded by the 16 NIH Institutes and Centers that support the NIH Blueprint for Neuroscience Research; and by the McDonnell Center for Systems Neuroscience at Washington University. We thank the reviewers whose comments/suggestions helped improve and clarify this manuscript. We also thank researchers processing Human Connectome Project data for providing the pre-processed data and rfMRI netmats.

## Code availability

Matlab scripts were written to perform the analyses described; this code is available from the authors upon request.

## Footnotes

1

2 https://www.humanconnectome.org/documentation/S1200/

3 The first eigenvector contains constant values.

4 http://www.matthewtoews.com/

5 DTI=FA+MD+RD+AD; rfMRI netmat= partial correlation and ICA-100

6 Results for full siblings are reported in Table 7 of Supplement material.

